# Species-specific imprinting of *Adam23* and its implications for gyrencephalic brain development

**DOI:** 10.1101/2025.12.01.691626

**Authors:** Makiko Meguro-Horike, Kengo Saito, Miyuki Shimazu, Yohei Shinmyo, Hiroshi Kawasaki, Shin-ichi Horike

**Affiliations:** Research Center for Experimental Modeling of Human Disease, Division of Integrated Omics research, Kanazawa University, Ishikawa, Japan; Department of Medical Neuroscience, Graduate School of Medical Sciences, Kanazawa University, Ishikawa, Japan; Department of Neurophysiology, Hamamatsu University School of Medicine, Shizuoka, Japan; Sapiens Life Sciences, Evolution and Medicine Research Center, Kanazawa University, Ishikawa, Japan

**Keywords:** Genomic imprinting, *Adam23*, Ferret brain, Neuronal differentiation, Brain evolution

## Abstract

Genomic imprinting is a parent-of-origin–specific epigenetic mechanism with essential roles in placental development and fetal growth in mice, but its contribution to brain function and evolution remains unclear. Here, we investigated genomic imprinting in the ferret, a gyrencephalic mammal whose cortical architecture closely resembles that of humans rather than mice. Guided by human imprinting datasets, we screened 65 candidate genes in ferret brain and identified paternal expression of *Adam23*, maternal expression of *Atp10a*, and biallelic expression of genes including *Pxdc1*, *Wrb* and *Ube3a*. *Adam23* showed paternal expression in ferret cortex, particularly in the occipital region, whereas it was biallelically expressed in mouse brain. In human SH-SY5Y cells, *ADAM23* was biallelic in undifferentiated cells but shifted toward monoallelic expression upon neuronal differentiation. CpG analysis of upstream islands indicated that *Adam23* imprinting in ferrets is independent of promoter methylation, suggesting a non-canonical mechanism. Functional assays in Neuro-2aTG cells showed that *Adam23* knockdown enhances neurite outgrowth, whereas overexpression promotes cell proliferation. These findings identify *Adam23* as a non-canonical imprinted gene whose dosage influences stem cell dynamics and neurite development, linking imprinting to the evolution of gyrencephalic cortical structures and highlighting the ferret as a valuable model for imprinting studies in higher-order brain function.

## Introduction

Genomic imprinting is an intriguing process that governs parent-of-origin-specific monoallelic expression of certain genes through differential epigenetic marks of the two parental alleles during germ cell development [1, 2]. This parent-of-origin-specific monoallelic expression results in a functional asymmetry between the maternal and paternal genomes, emphasizing the crucial requirement for contributions from both parental genomes for proper development [3]. Thus, genomic imprinting is undoubtedly a fascinating departure from classical Mendelian genetics, and its study has given rise to numerous hypotheses concerning its biological and evolutionary significance in mammals [4, 5]. These hypotheses, including the conflict hypothesis, prevention-of-parthenogenesis theory, genome protection from foreign genes, and the placental hypothesis, provide different angles for understanding why genomic imprinting exists and what roles it plays in mammalian biology[6–10]. Moreover, the variation in genomic imprinting among mammalian species further complicates the picture. To date, more than a few hundred imprinted genes have been identified, and although many are conserved across mammalian species, some display significant divergence in their imprinting patterns [11–14]. These interspecies differences are not only important for understanding the evolutionary significance of imprinting but also underscore the limitations of modeling human imprinting-related neurological disorders in animals. For example, abnormalities in the 15q11–q13 region underlie the neurodevelopmental disorders Prader–Willi syndrome (PWS) and Angelman syndrome (AS)[15–17]. Mouse models of these disorders, although valuable, do not fully capture the complexity of the human phenotypes [18–20]. Similarly, duplications of the same 15q11–q13 region, which are associated with autism, also remain poorly understood, largely because parental allele-specific effects differ between humans and mice [21]. Therefore, there is growing interest in non-human primate models, such as marmosets and rhesus monkeys, which share greater genetic similarity with humans and may provide more faithful insights into the role of imprinting in neurological and psychiatric disorders[22, 23]. However, progress has been hampered by the lack of informative genetic markers, such as SNPs, especially in non-inbred species, which remains a significant obstacle to advancing this field.

In this study, the ferret, which, unlike the mouse or marmoset, possesses a gyrencephalic brain structure more similar to that of humans, was chosen as an alternative model [24, 25]. This choice provided valuable insight into the role of genomic imprinting in higher brain function and structure. The identification of the *Adam23* gene, which is bi-allelically expressed in mice but paternally expressed in ferrets and humans, along with the positive effect of the *Adam23* gene on cell proliferation, provides intriguing insights into how genomic imprinting affects brain development. Furthermore, the evolutionary acquisition of genomic imprinting of the *Adam23* gene in ferrets and humans suggests its involvement in promoting neurite outgrowth, thereby contributing to the complexity of neural networks.

In summary, this study sheds light on the multifaceted world of genomic imprinting, its role in brain evolution, and its impact on neural functions. It underscores the need for diverse model systems and approaches to unravel the intricate mechanisms governing mammalian development and higher brain structures.

## Material and methods

### Ethics statement

All animal experiments were conducted in accordance with the institutional guidelines for the care and use of laboratory animals and were approved by the Animal Experiment Committee of Kanazawa University (Approval Nos. AP23-050-02, AP22-035-01 and AP24-071-02). All efforts were made to minimize animal suffering and to reduce the number of animals used.

### Animals

Ferrets (*Mustela putorius furo*) were purchased from Japan SLC (Hamamatsu, Japan) and Marshall Farms (North Rose, NY). Ferrets were housed as previously described [26], under a 16/8 h light/dark cycle with ad libitum access to food and water, at 20–22 °C and 45–65% relative humidity. C57BL/6J mice were purchased from Japan SLC, and JF1/Ms mice were provided by the Mouse Development Laboratory, National Institute of Genetics (Mishima, Japan)[27, 28]. Mice were maintained under a 12/12 h light/dark cycle with ad libitum access to food and water, at 20–26 °C and 45–65% relative humidity.

### Collection and preservation of brain tissue samples

Mice and ferrets were deeply anesthetized using a triple-anesthetic cocktail consisting of medetomidine (0.3 mg/kg), midazolam (4 mg/kg), and butorphanol (5 mg/kg), after which the brains were rapidly dissected without perfusion fixation. Dissected tissues were immediately snap-frozen in liquid nitrogen to preserve RNA integrity.

### Cells

Neuro-2aTG, a 6-thioguanine-resistant (HAT-sensitive mutant) variant of Neuro-2a cells (JCRB cell bank, Osaka, Japan)[29], was cultured in Dulbecco’s Modified Eagle Medium (DMEM; Wako, Osaka, Japan) supplemented with 10% fetal bovine serum (FBS; Gibco, Thermo Fisher Scientific, USA) and 100 µg/ml penicillin-streptomycin (Wako). Cells were maintained at 37 °C in a humidified atmosphere of 5% CO₂.

SH-SY5Y, a human neuroblastoma cell line (provided by Dr. Janine LaSalle, University of California, Davis, USA)[30], was cultured in DMEM/F12 (Wako) supplemented with 15% fetal bovine serum (FBS; Gibco, Thermo Fisher Scientific, USA) and 100 µg/ml penicillin-streptomycin (Wako). Cells were maintained at 37 °C in a humidified atmosphere of 5% CO₂.

### DNA Extraction

Genomic DNA was isolated from cultured cells and tissue fragments using the Gentra Puregene Cell Kit (QIAGEN, Hilden, Germany) according to the manufacturer’s instructions. DNA concentration and purity were assessed with a NanoDrop ND-1000 spectrophotometer (Thermo Fisher Scientific, USA).

### RNA Extraction and cDNA Synthesis

Total RNA was extracted from cultured cells and tissue fragments using the RNeasy Mini Kit (QIAGEN, Hilden, Germany) according to the manufacturer’s instructions, including on-column DN*ase* I digestion (QIAGEN) to remove genomic DNA contamination. RNA concentration and purity were determined with a NanoDrop ND-1000 spectrophotometer (Thermo Fisher Scientific, USA). First-strand cDNA was synthesized from total RNA using SuperScript™ III Reverse Transcriptase (Thermo Fisher Scientific, USA) with random hexamer primers (Invitrogen, USA) at 42 °C for 30 min.

### PCR and Sequencing Analysis

Single nucleotide polymorphism (SNP) and gene expression analyses were performed by PCR using GoTaq® G2 Hot Start Green Master Mix (Promega, USA). The PCR conditions were as follows: 95 °C for 2 min; 35 cycles of 95 °C for 30 s, 58 °C for 30 s, and 72 °C for 30 s; followed by a final extension at 72 °C for 5 min. PCR products were verified by electrophoresis on 2% agarose gels.

PCR products were treated with Shrimp Alkaline Phosphatase (Thermo Fisher Scientific, USA) and Exonuclease I (New England Biolabs, USA) and subsequently purified prior to sequencing. Sequencing was performed using the SupreDye™ Cycle Sequencing Kit (MS Techno Systems, Japan) according to the manufacturer’s instructions. Sequencing data were analyzed with 4Peaks software (https://nucleobytes.com/4peaks/).

The primer sequences used for SNP analysis are listed in Table 1. For determination of the gap sequence upstream of the ferret *Adam23* gene, the following primers were used: fAdam23_GAP_F1, 5′-AGC ATC AGC ATC AGC ATC AG-3′; fAdam23_GAP_R1, 5′-TGC AAC AGA AAT GGA CTT CA-3′.

**Table 1.**
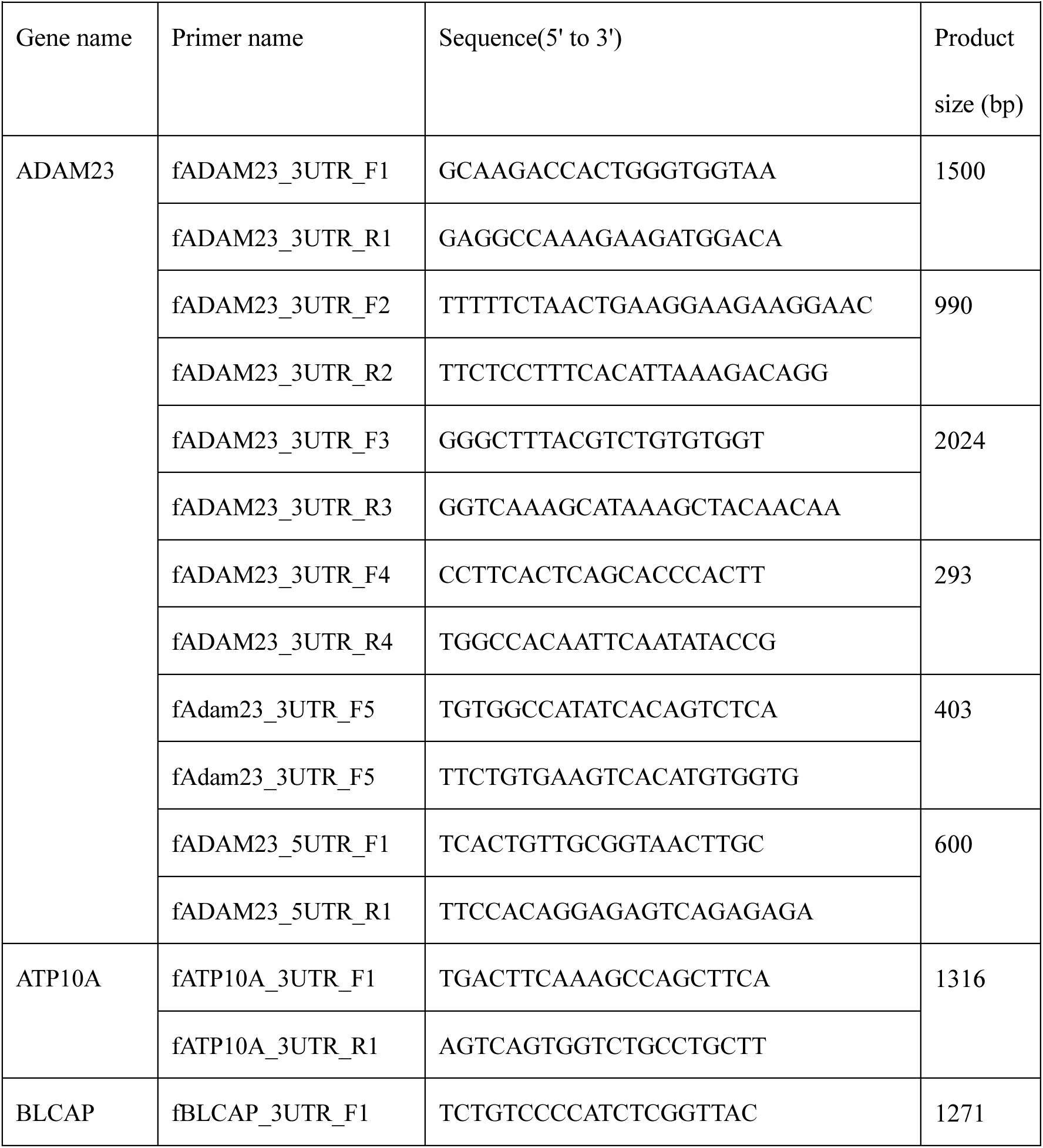
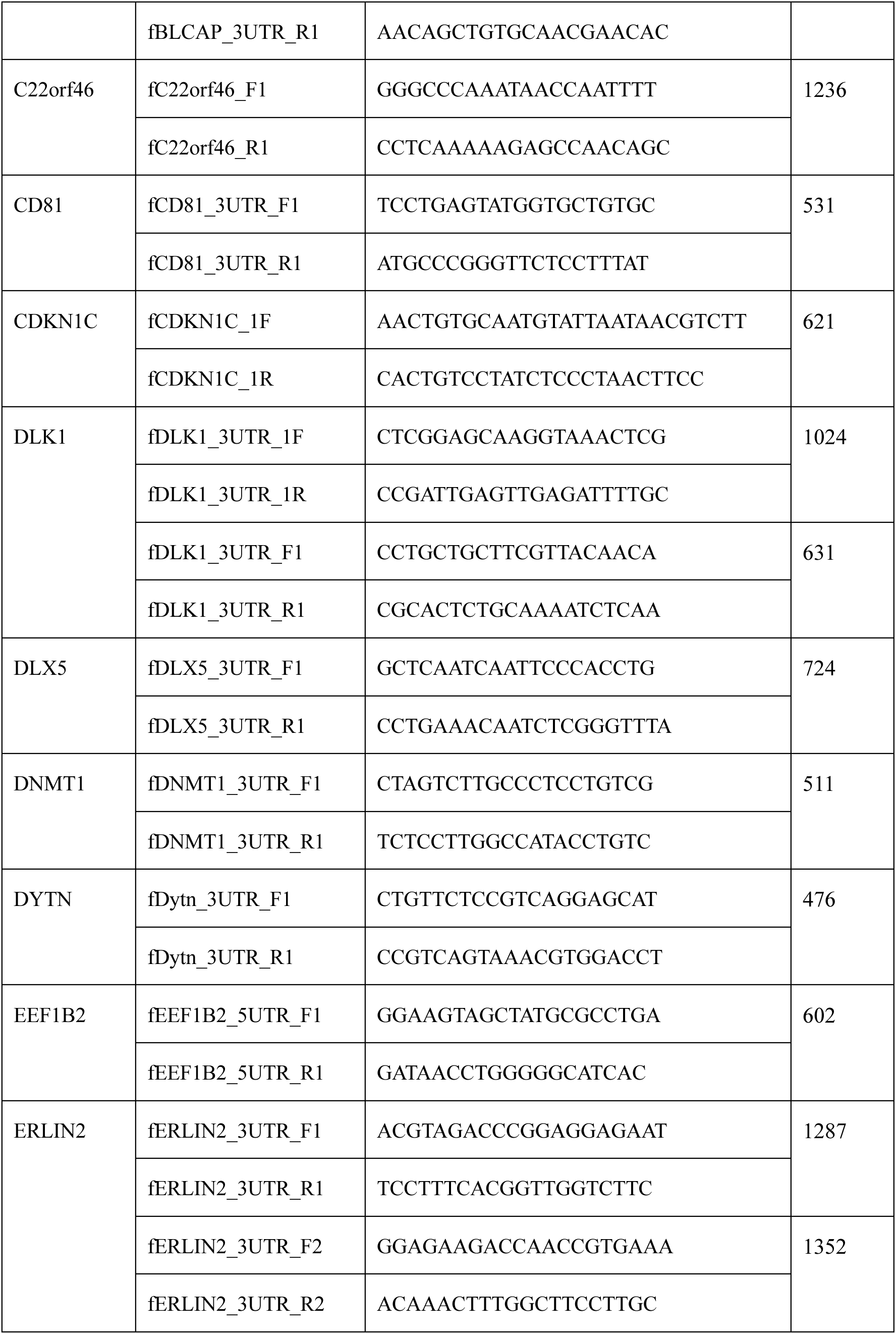
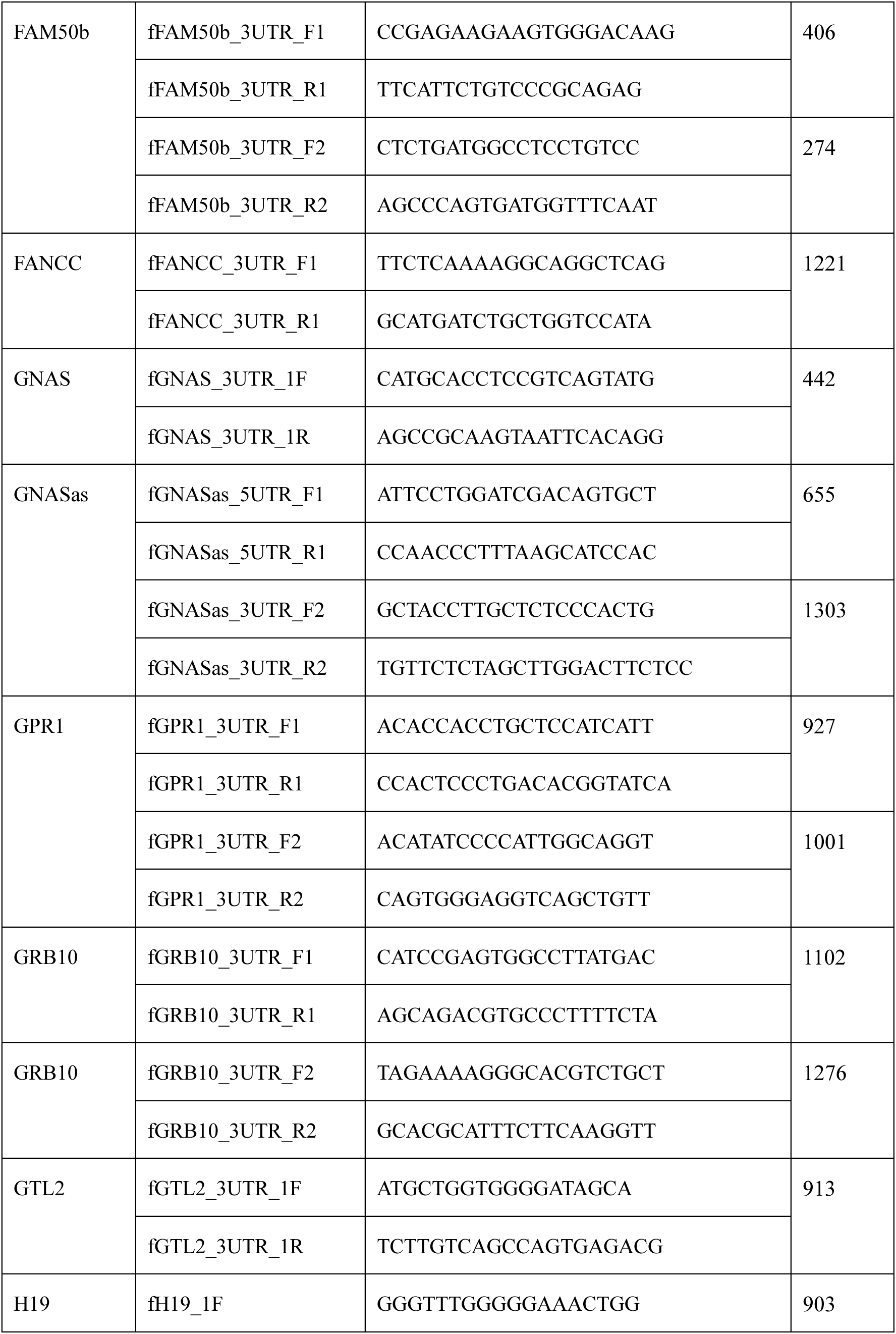
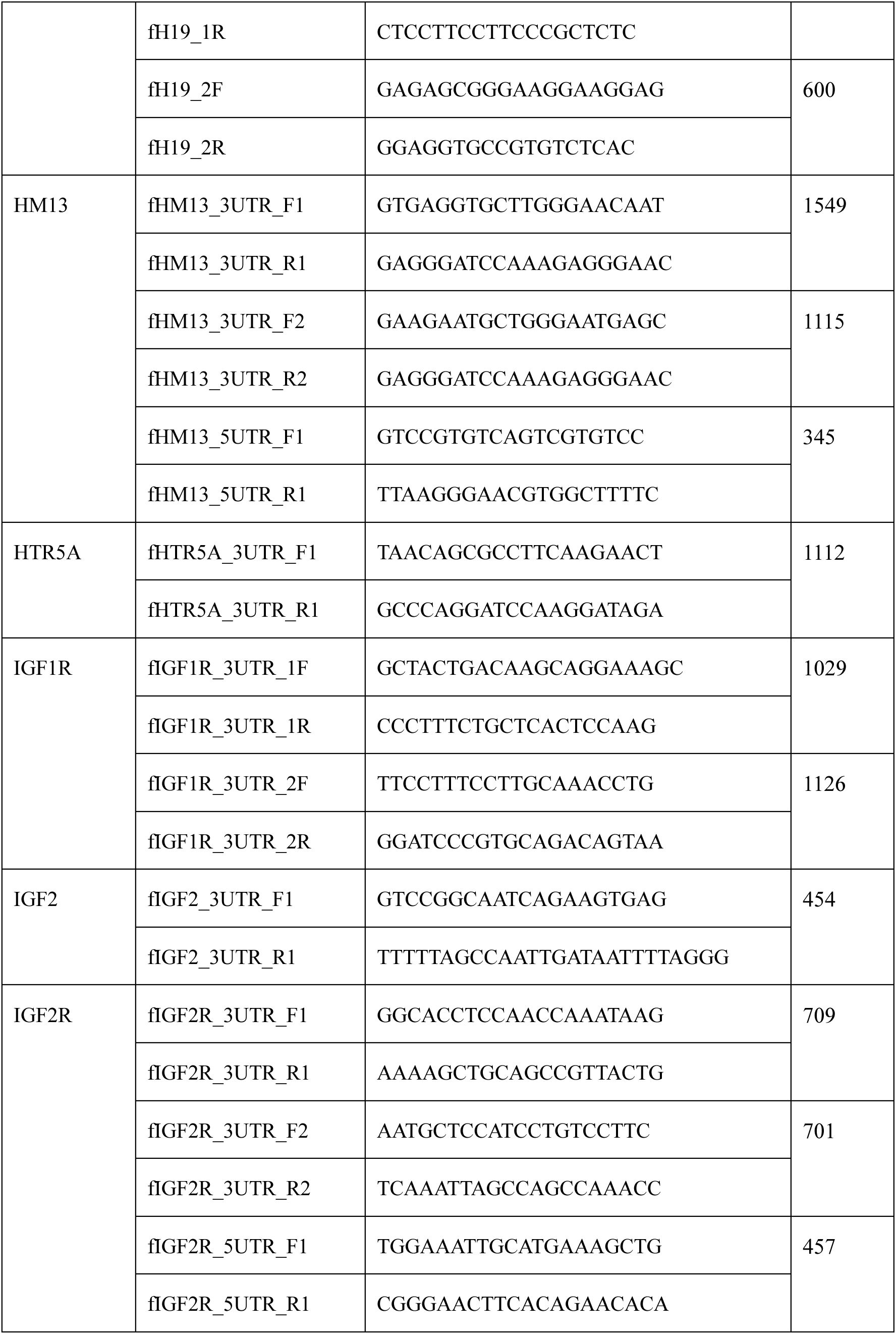
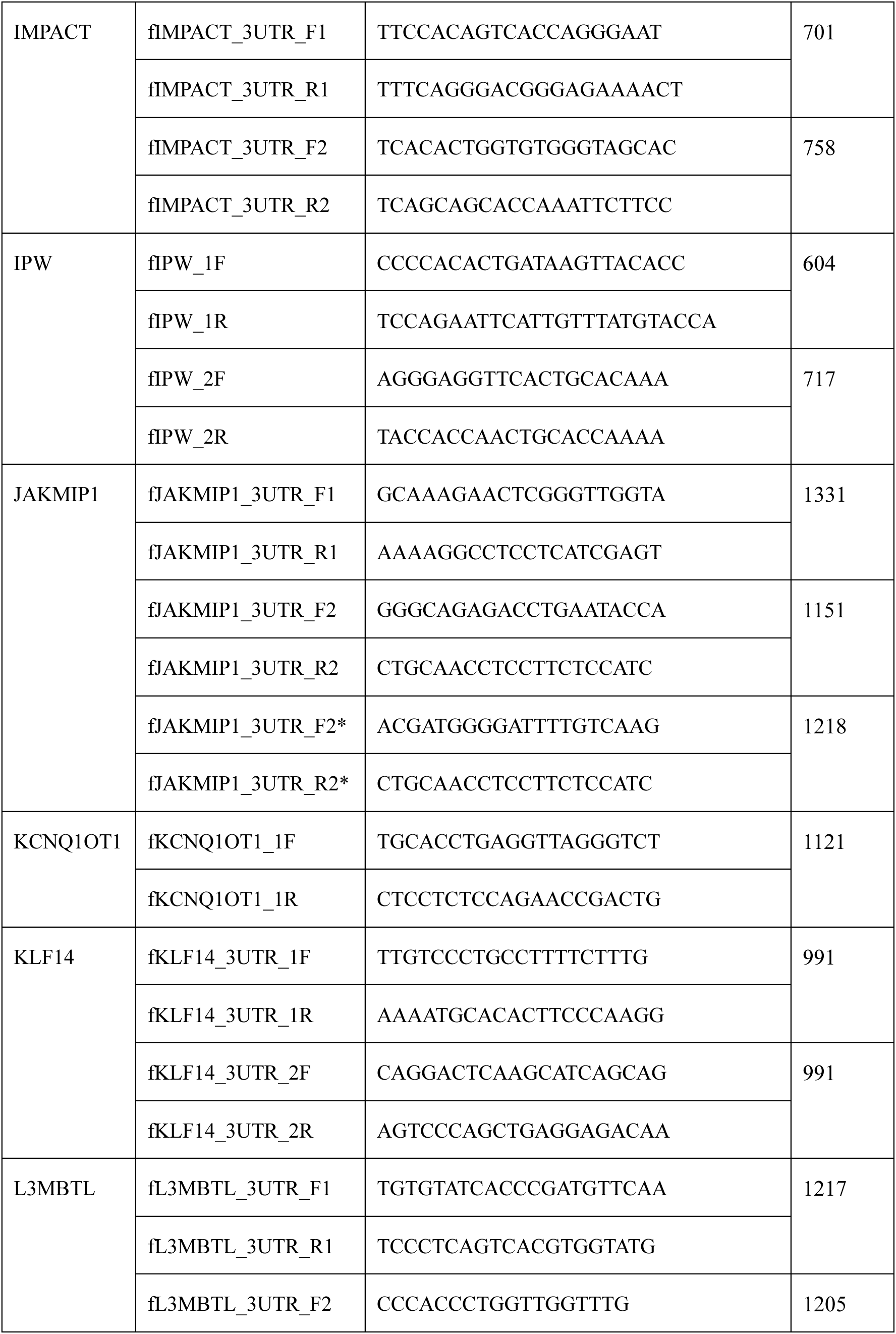
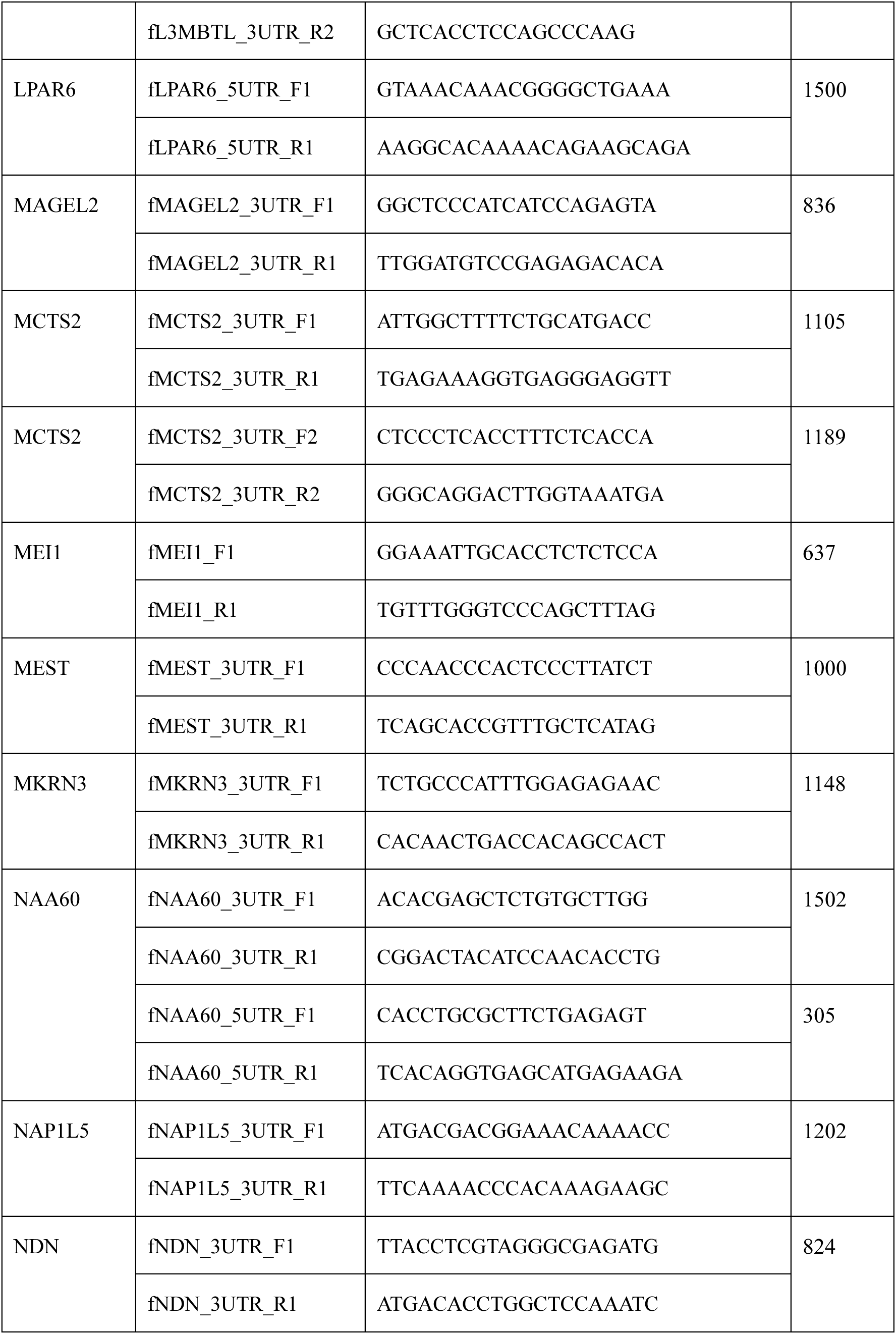
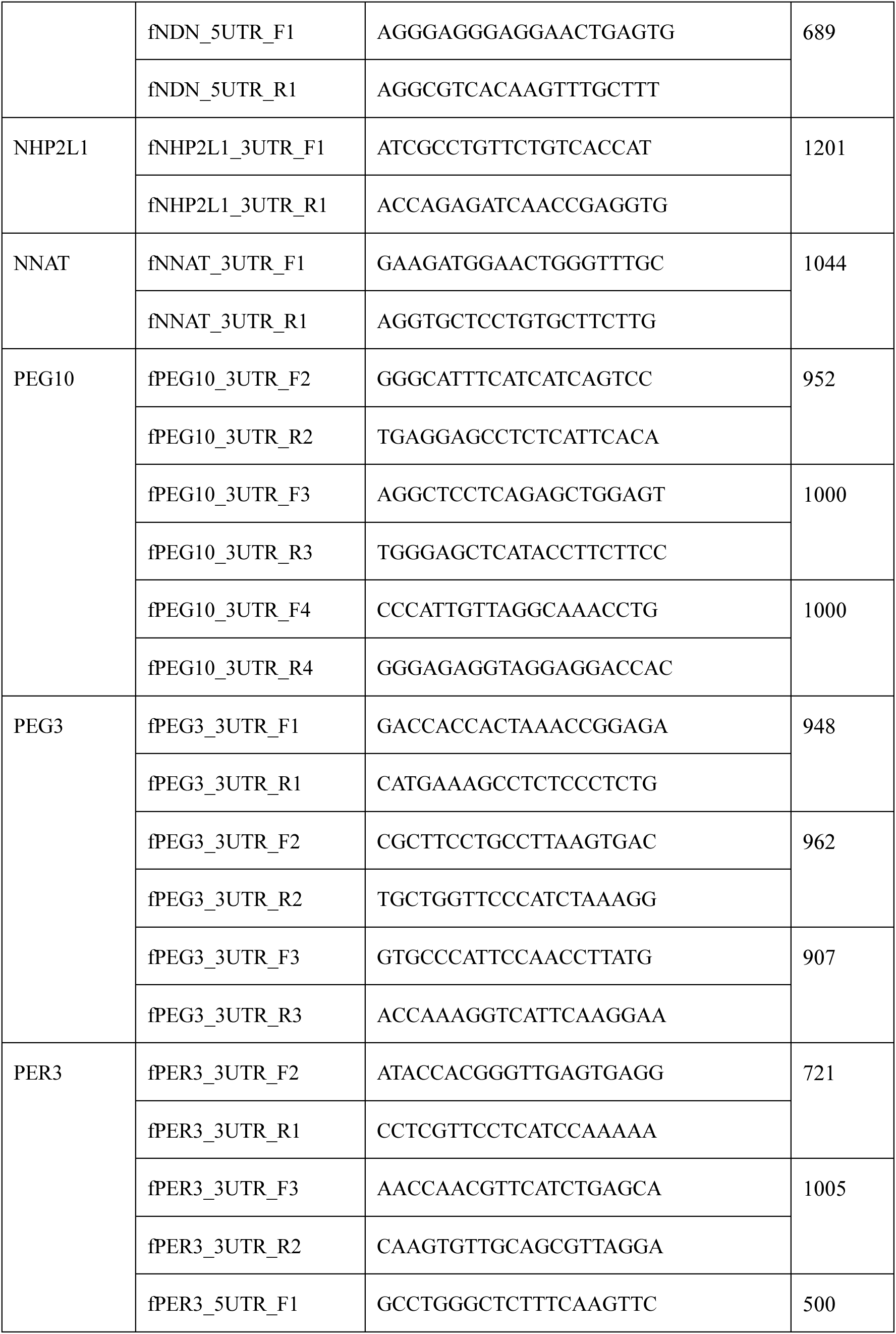
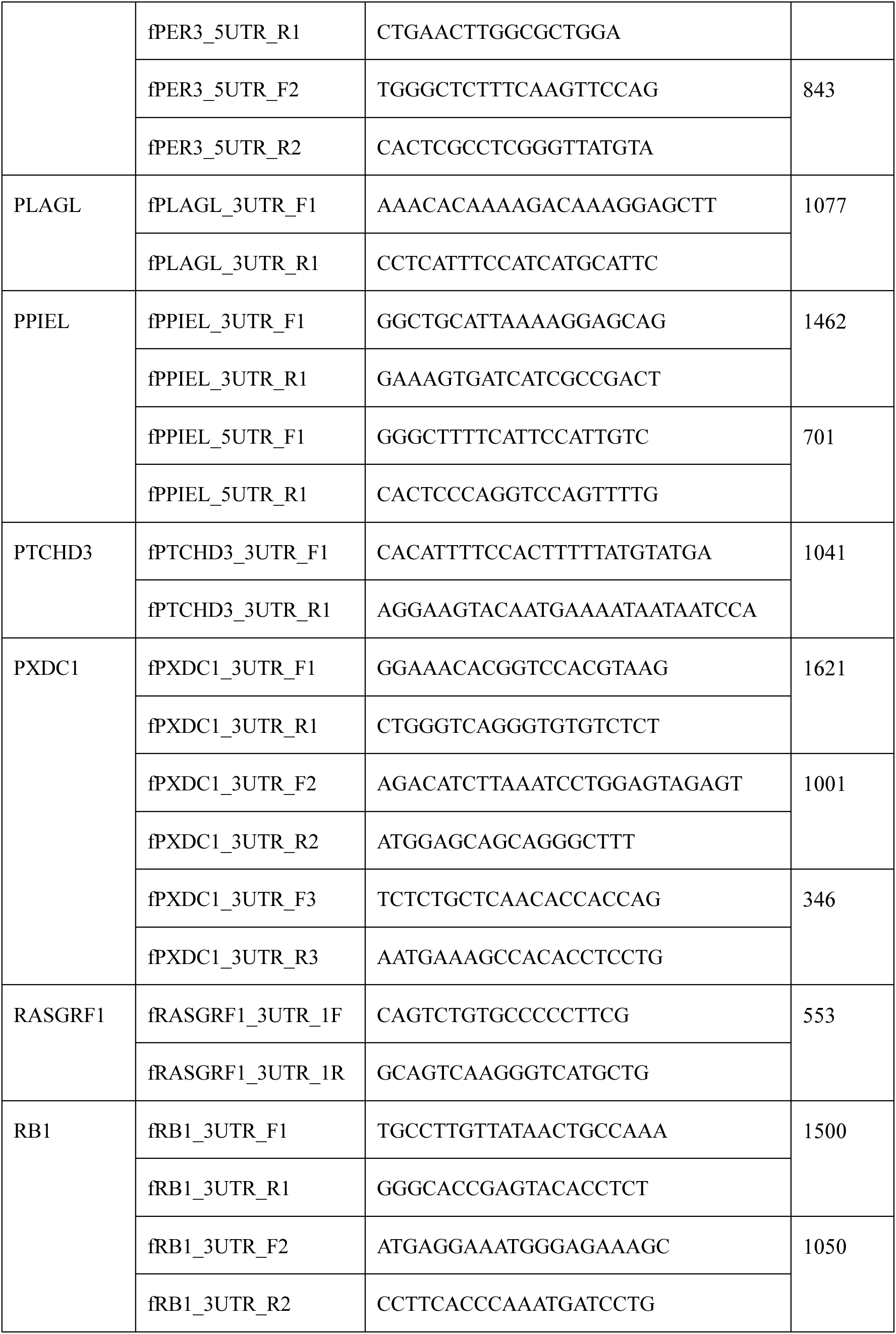
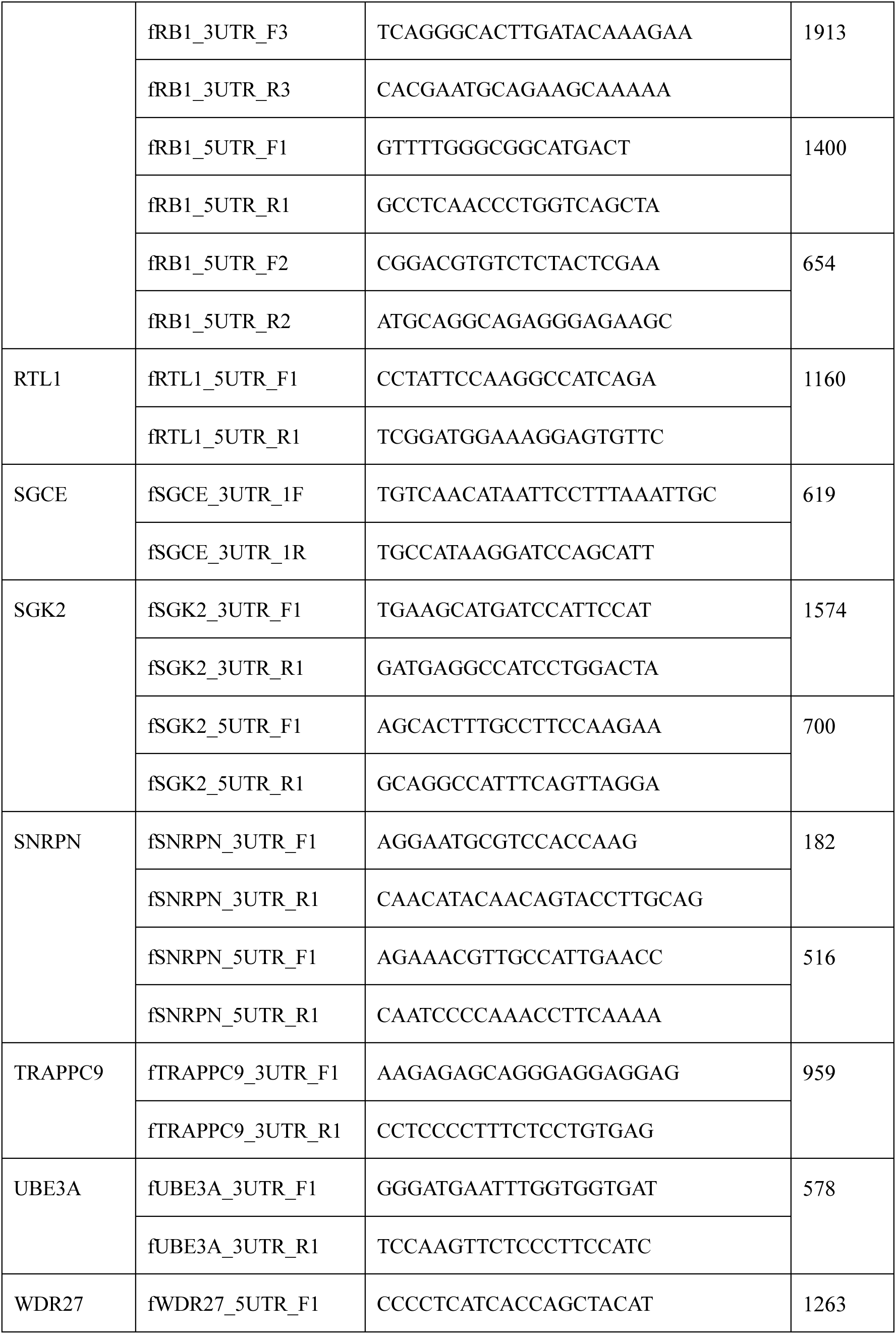
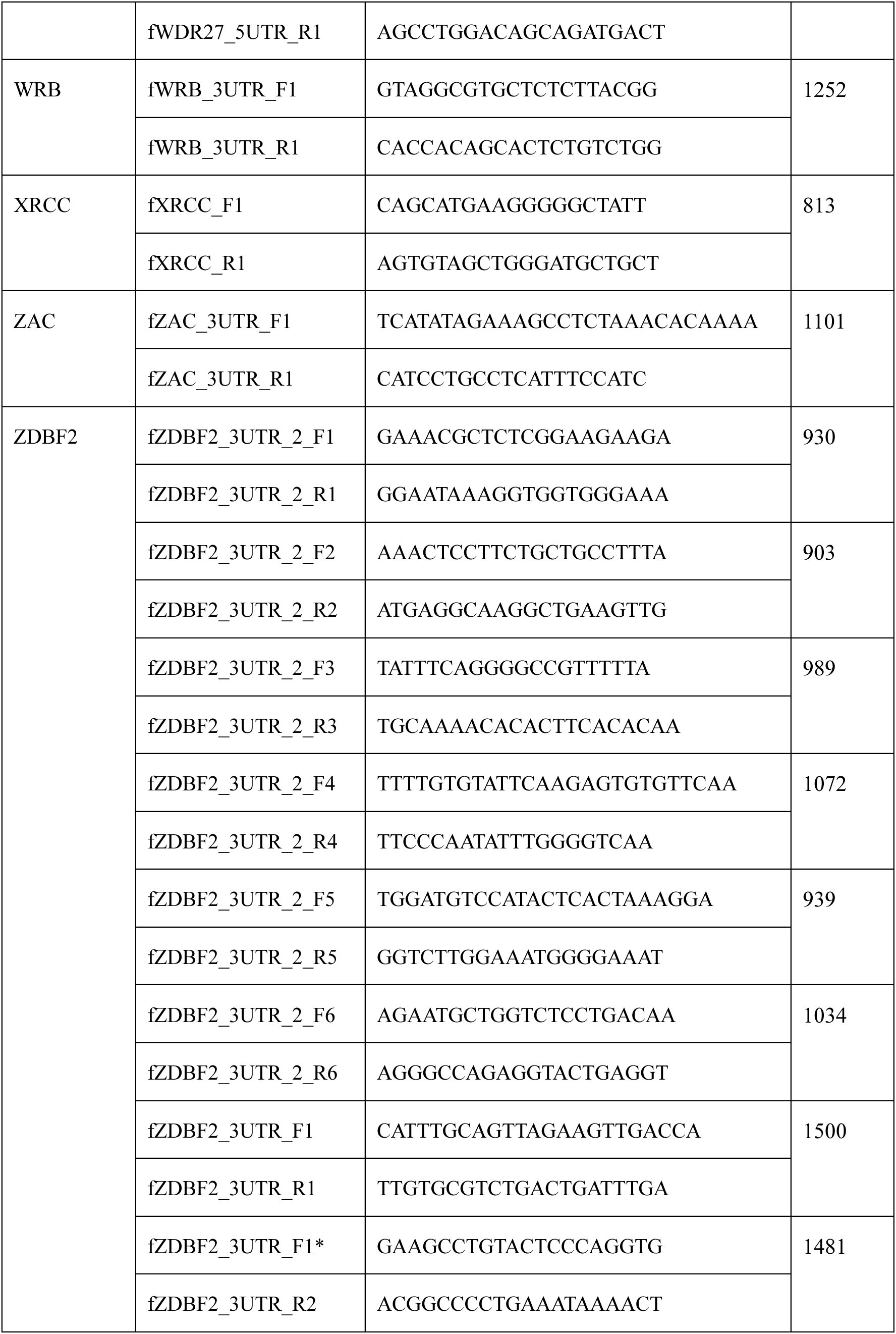
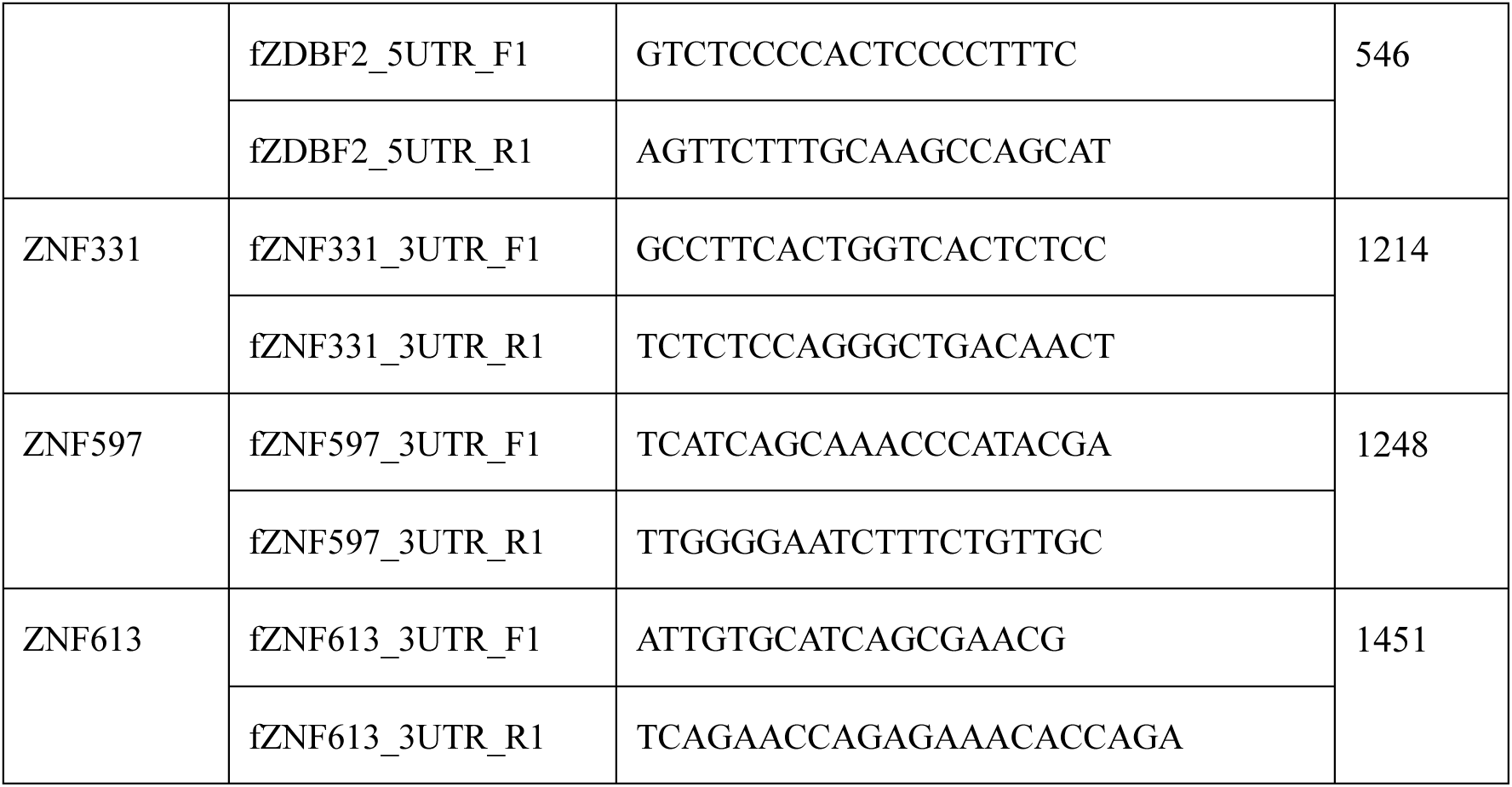
List of 65 candidate genes analyzed for genomic imprinting in the ferret and the primer sequences used for their analysis.

### DNA Methylation Analysis

DNA methylation analysis was performed using the EZ DNA Methylation-Gold Kit (Zymo Research, USA) according to the manufacturer’s protocol. Bisulfite-treated DNA was used as a template for PCR amplification of four segments (I–IV) covering the upstream CpG island region of the ferret *Adam23* gene. Primer sequences were designed with MethPrimer (https://www.urogene.org/cgi-bin/methprimer/methprimer.cgi) and are listed below:

CpG I: fAdam23_CpG_F1, 5′-TGT TTG GTA ATA GTA ATA GTA TTA GTA TTA-3′; fAdam23_CpG_R1, 5′-CTT AAA ACC TCC CCA AAT ACT CC-3′

CpG II: fAdam23_CpG_F2, 5′-GGA GGG AGG AGT ATT TGG GG-3′; fAdam23_CpG_R2, 5′-ACT TCA TAA CTC ATA ACT CCC C-3′

CpG III: fAdam23_CpG_F3, 5′-GGG GGA GTT ATG AGT TAT GAA GT-3′; fAdam23_CpG_R3, 5′-CTA AAT ACA ACA ATC CCC CAA AC-3′

CpG IV: fAdam23_CpG_F4, 5′-TGA AGG TTA AGT AGG AGG TTT TGT T-3′; fAdam23_CpG_R4, 5′-AAA TTC TTC CCC AAA TTC TTC C-3′

PCR products were TA-cloned using the pGEM™-T Easy Vector System (Promega, USA) and sequenced with T7 or SP6 primers. At least ten independent clones per sample were sequenced to determine methylation status.

### Knockdown Experiment of Mouse *Adam23* Gene in Neuro-2aTG Cells

#### A) Establishment of Mouse *Adam23* Knockdown Cells

Neuro-2aTG cells were transfected with the MISSION shRNA vector targeting mouse *Adam23* (Sigma-Aldrich, Clone ID: TRCN0000032662) using Lipofectamine™ 2000 (Thermo Fisher Scientific, USA). After 48 h, cells were subjected to puromycin selection (2 µg/ml) to establish stable *Adam23* knockdown clones. Control cells were generated by transfection with the MISSION non-target shRNA Control Vector (Sigma-Aldrich, SHC016) under the same conditions. Three independent *Adam23* knockdown clones, as well as three independent control clones, were established and subsequently used for all analyses.

#### B) Assessment of Knockdown by Real-Time PCR

Total RNA was extracted, and *Adam23* transcript levels were quantified by real-time PCR using SsoAdvanced Universal SYBR Green Supermix (Bio-Rad, USA). *Gapdh* was used as an internal reference, and relative expression levels were calculated by the ΔΔCt method. The primers used were:

Adam23: mAdam23_qF1, 5′-CGG CTG CAT CAT GGA AGA AA-3′; mAdam23_qR1, 5′-TTG AAA AGA CAT GCT CCG CC-3′

Gapdh: mGapdh_qF1, 5′-ACC ACA GTC CAT GCC ATC AC-3′; mGapdh_qR1, 5′-TCC ACC ACC CTG TTG CTG TA-3′

Of the three established knockdown clones, two were selected for analysis of *Adam23* knockdown efficiency.

### Experimental Overexpression of Bovine *Adam23* Gene in Neuro-2aTG Cells

#### A) Establishment of Cells Overexpressing Bovine *Adam23* Gene

Neuro-2aTG cells were transfected with a vector carrying bovine *Adam23* cDNA (Horizon Discovery, Clone ID: 8208020) under the control of the CAG promoter using Lipofectamine™ 2000 (Thermo Fisher Scientific, USA). After 48 h, cells were subjected to G418 selection (400 µg/ml) to establish stable *Adam23*-overexpressing clones. Control cells were generated by transfection with an empty vector lacking cDNA downstream of the CAG promoter. Three independent *Adam23*-overexpressing clones, as well as three independent control clones, were established and subsequently used for all analyses.

#### B) RT-PCR Specific for Bovine *Adam23* Gene

Overexpression of bovine *Adam23* was confirmed by RT-PCR using GoTaq® G2 Hot Start Green Master Mix (Promega, USA). PCR products were visualized on 2% agarose gels. The following primers were used:

Bovine *Adam23*: bAdam23_qF1, 5′-CAA TGG TTT GCT GTC TTC TGA-3′; bAdam23_qR1, 5′-CCG GTC ACT GCT TTC CAT AC-3′

Mouse *Gapdh:* mGapdh_qF1, 5′-ACC ACA GTC CAT GCC ATC AC-3′; mGapdh_qR1, 5′-TCC ACC ACC CTG TTG CTG TA-3′

### Neurodifferentiation Experiment of Neuro-2aTG Cells

Neuro-2aTG cells were seeded in Lab-Tek™ II Chamber Slides (Nunc, USA) at a density of 1 × 10⁵ cells per well. Cells were cultured in DMEM (Wako, Japan) supplemented with 0.1% fetal bovine serum (FBS; Gibco, Thermo Fisher Scientific, USA) and 100 µg/ml penicillin-streptomycin (Wako). Neurodifferentiation was induced by treatment with 20 µM retinoic acid (Wako) for 3 days at 37 °C in a humidified 5% CO₂ incubator.

### Neuronal Differentiation of SH-SY5Y Cells

SH-SY5Y cells were seeded at a density of 3 × 10⁵ cells per 35-mm dish and cultured in DMEM/F12 (Wako, Japan) supplemented with 15% fetal bovine serum (FBS; Gibco, Thermo Fisher Scientific, USA), 2 mM L-glutamine (Wako), and 100 µg/ml penicillin-streptomycin (Wako) at 37 °C in a humidified atmosphere of 5% CO₂.

Neuronal differentiation was induced over an 18-day protocol involving sequential changes in culture medium. From day 1 to day 7, cells were maintained in DMEM/F12 supplemented with 2.5% FBS, 2 mM L-glutamine, 10 µM retinoic acid, and antibiotics. From day 8 to day 10, cells were cultured in DMEM/F12 containing 1% FBS, 2 mM L-glutamine, 10 µM retinoic acid, and antibiotics on ECM-coated dishes. From day 11 to day 18, cells were cultured in Neurobasal™ medium (Gibco) supplemented with B27 (1×), 20 mM KCl, 2 mM GlutaMAX™ (Gibco), 50 ng/ml BDNF (Wako), 2 mM dibutyryl cyclic AMP (db-cAMP; Functional Peptide Institute), 10 µM retinoic acid, and antibiotics, with medium changes every 3–4 days.

On day 18, RNA was extracted from differentiated cells for gene expression analysis.

### Cell Proliferation Assay

Neuro-2aTG cells were seeded in 6-well plates (Nunc, USA) at a density of 5 × 10⁴ cells per well. Cell numbers were counted daily for 5 consecutive days starting from day 0 (the day after seeding). Relative cell counts were calculated with the value at day 0 set as 1.

### Immunostaining

Neuro-2aTG cells cultured on Lab-Tek™ II chamber slides (Nunc, USA) were fixed with 4% paraformaldehyde (PFA) in PBS and permeabilized with 0.1% Triton X-100/PBS. After blocking (Blocking Reagent, Roche), cells were incubated with mouse anti-βIII tubulin antibody (Merck) overnight at 4 °C, followed by Cy3-conjugated secondary antibody (Merck) for 1 h at 37 °C. Nuclei were counterstained with DAPI and mounted in VECTASHIELD Antifade Mounting Medium (Vector Laboratories, USA).

Fluorescence images were acquired using an Olympus fluorescence microscope equipped with a Qimaging CCD camera. Neurite length was quantified from βIII tubulin-stained cells using iVision software (BioVision Technologies, USA).

### Statistical Analysis

Data are presented as mean ± standard error (SE). Statistical analyses were performed using Prism 5.0 software (GraphPad Software, USA). Neurite outgrowth was compared using the Mann–Whitney U test. Cell proliferation curves were analyzed by two-way analysis of variance (ANOVA) followed by Bonferroni’s multiple comparison test. A p-value < 0.05 was considered statistically significant.

## Results

### Search for Imprinted Genes in the Ferret Brain

To investigate genomic imprinting in the ferret brain, we first curated a set of candidate genes previously reported as imprinted in humans. Curt et al. identified parental allele–specific differentially methylated regions across blood, brain, and liver using genome-wide methylation arrays [31], whereas Jadhav et al. defined parent-of-origin–dependent transcription using RNA-seq in human blood [32]. Guided by these datasets, we selected 65 genes for imprinting analysis in the ferret (Supplementary Figure 1a, Table 1). To establish whether these genes harbored informative variants suitable for assigning parental origin, we amplified and sequenced the 5′ and 3′ untranslated regions (UTRs) of each gene using genomic DNA from ferret trios (father–mother–offspring). This screen identified single nucleotide polymorphisms (SNPs) in eight genes (*Adam23*, *Atp10a*, *Fam50b*, *Nhp2l1*, *Ptchd3*, *Pxdc1*, *Ube3a*, and *Wrb*) (Supplementary Figure 1a). *Fam50b* and *Ptchd3* were not detectably expressed in the ferret brain and were therefore excluded from further examination (Supplementary Figure 2). The remaining six genes were subjected to allele-specific expression analysis. Allelic ratios were quantified from chromatogram peak heights at informative SNP positions (Supplementary Figure 1b). In accordance with the canonical 70:30 threshold used in previous imprinting studies [33–36], paternal or maternal imprinting was defined as an allele ratio >2.33 or <0.43, respectively. *Pxdc1* and *Wrb* (Supplementary Figure 3a, b), as well as *Ube3a* (Supplementary Figure 3c), exhibited biallelic expression (0.78–0.96), diverging from human reports—a discrepancy that may partly reflect differences in the tissues examined. In contrast, *Atp10a* displayed maternal bias (0.37–0.52) (Supplementary Figure 4), while *Adam23* (Figure 1b) and *Nhp2l1* (data not shown; full report in preparation) exhibited unequivocal paternal expression.

**Figure 1.**
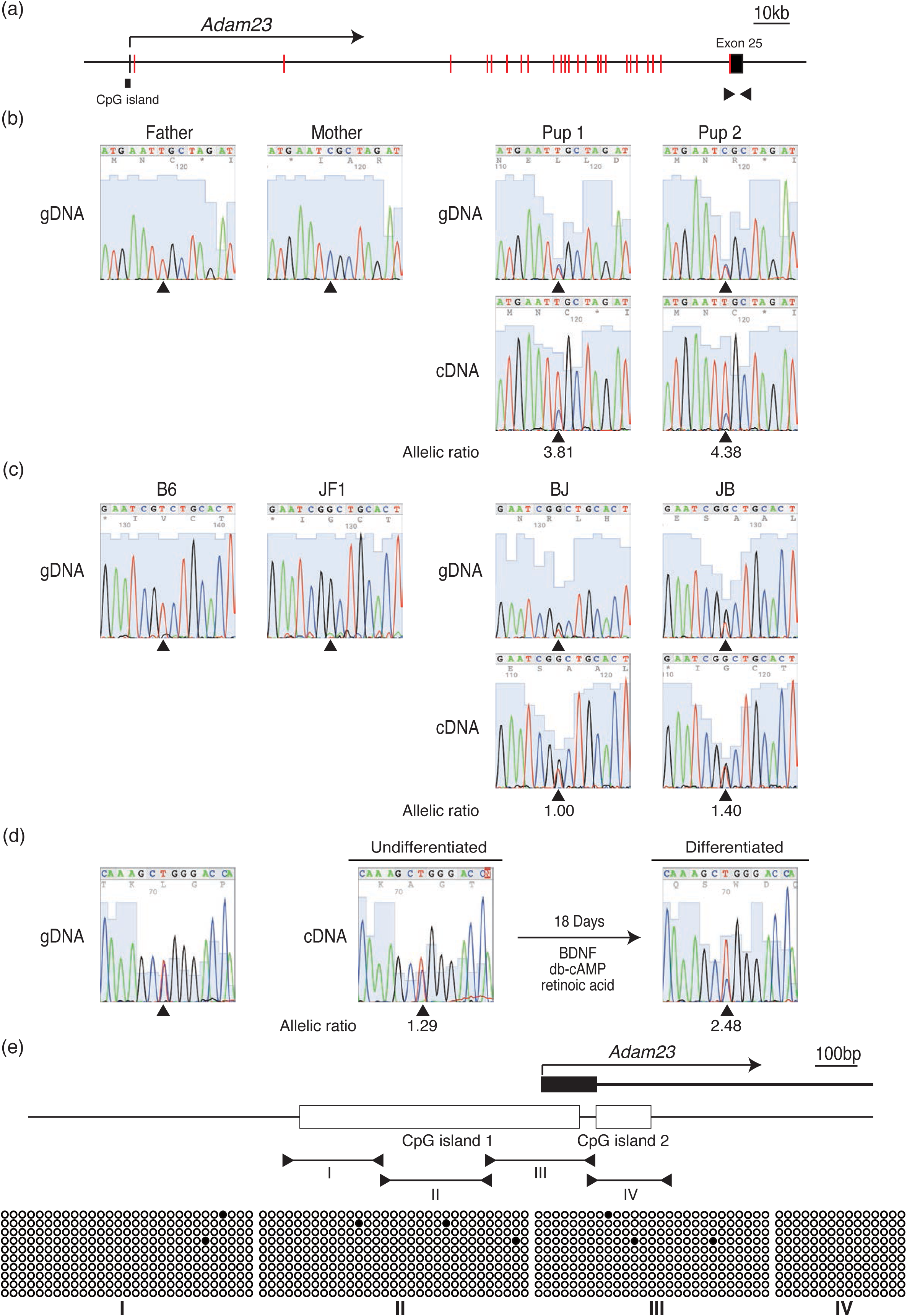
Ferret *Adam23* exhibits paternal expression. (a) Genomic structure of the ferret *Adam23* locus (GL897001: 5,240,001–5,440,000). Coding sequences (CDS) are shown in red, and untranslated regions (UTRs) are shown in black. Arrowheads indicate primer positions used for SNP screening. (b) Imprinting analysis showed that the ferret *Adam23* gene exhibits paternal expression. Arrowheads indicate polymorphic sites used for imprinting analysis. Allelic ratios calculated for imprinting analysis are shown beneath the cDNA chromatograms. gDNA, genomic DNA; cDNA, complementary DNA. Pup1 and Pup2 represent offspring derived from the parental combinations shown on the left (Father × Mother). (c) The mouse *Adam23* gene displays biallelic expression in brain tissues from reciprocal crosses between B6 and JF1 mice. Arrowheads indicate polymorphic sites used for imprinting analysis. Allelic ratios are shown beneath the cDNA chromatograms. BJ indicates offspring derived from B6 females crossed with JF1 males, whereas JB indicates offspring derived from JF1 females crossed with B6 males. (d) Changes in monoallelic expression of the human *ADAM23* gene during neuronal differentiation of SH-SY5Y cells. Arrowheads indicate polymorphic sites used for imprinting analysis. Allelic ratios are shown beneath the cDNA chromatograms. (e) DNA methylation analysis of the CpG island upstream of the ferret *Adam23* gene. Bisulfite sequencing was performed using primer sets I–IV. Open circles indicate unmethylated CpGs, and filled circles indicate methylated CpGs.

### Detailed Characterization of *Adam23* Imprinting in the Ferret Brain

Because *Adam23* showed robust paternal expression in our initial screen and has been implicated in higher-order neural functions, particularly synaptic connectivity[37–39], we next undertook a detailed characterization of its imprinting status. A C/T SNP in the 3′ UTR of exon 25 (Figure 1a) provided an unambiguous marker for distinguishing parental alleles: the father was homozygous T/T and the mother C/C.

RT–PCR followed by direct sequencing confirmed strong paternal expression of *Adam23* in the occipital cortex (allele ratio 3.81–4.38; Figure 1b). To determine whether imprinting varied across brain regions, we assessed allele-specific expression in the olfactory bulb, frontal cortex, parietal cortex, and occipital cortex. A pronounced paternal bias was observed in the occipital cortex, whereas the olfactory bulb displayed a near-biallelic ratio (1.7), approaching allelic equilibrium (Figure 2). These region-dependent differences indicate that *Adam23* imprinting is modulated in a context-specific manner and may contribute to functional specialization across gyrencephalic forebrain circuits.

**Figure 2.**
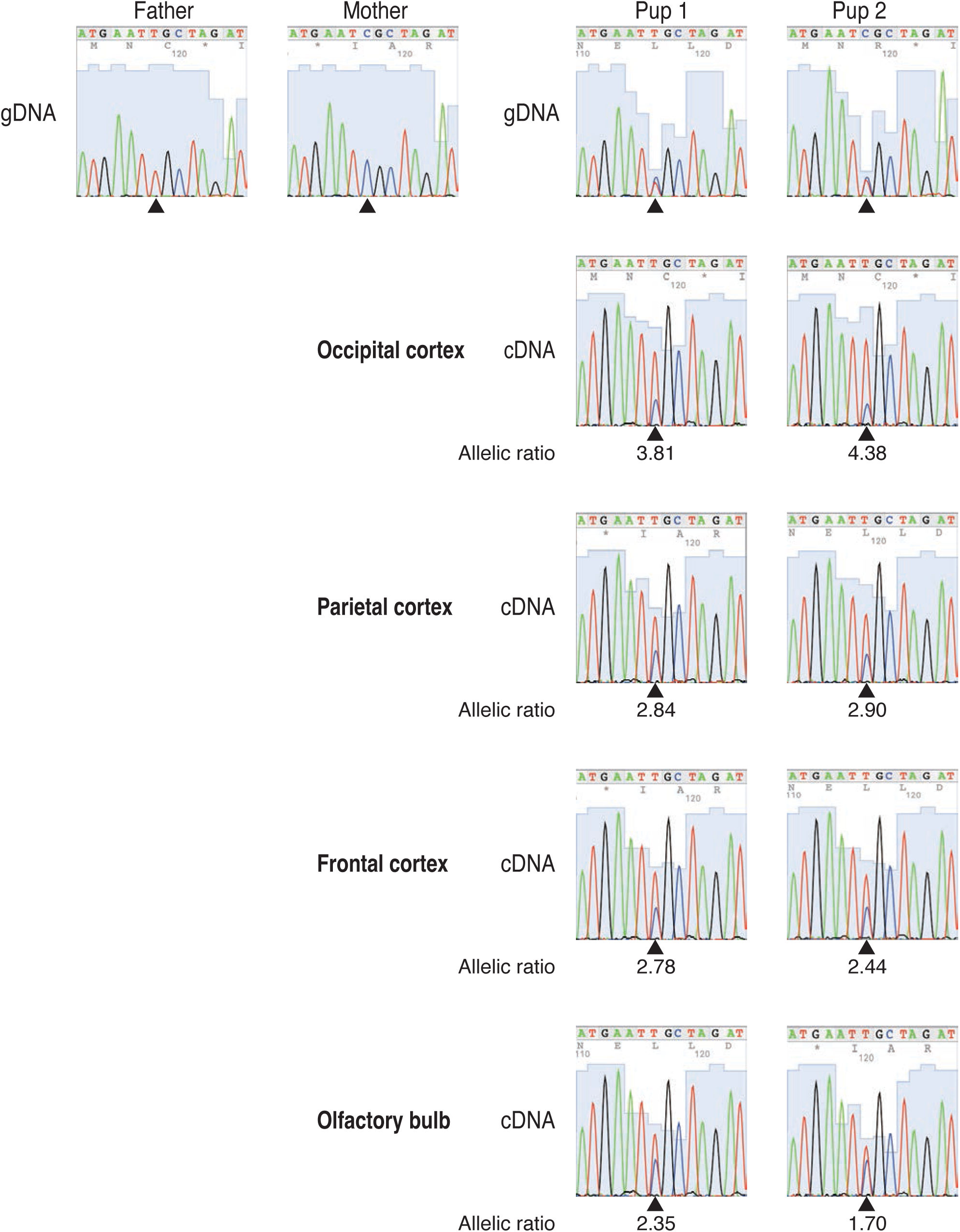
Paternal expression of *Adam23* in ferret brain tissues. Allele-specific expression of the *Adam23* gene in various ferret brain regions, including the olfactory bulb, frontal cortex, parietal cortex, and occipital cortex. Arrowheads indicate polymorphic sites used for imprinting analysis. Allelic ratios calculated for imprinting analysis are shown beneath the cDNA chromatograms. gDNA, genomic DNA; cDNA, complementary DNA. Pup1 and Pup2 represent offspring derived from the parental combinations shown on the left (Father × Mother).

### Species-Specific Genomic Imprinting of the *Adam23* Gene

Given that *Adam23* shows paternal expression in ferrets, we next asked whether this imprinting pattern is conserved across species. To examine the imprinting status of the *Adam23* gene in mice, we performed RT-PCR using RNA extracted from brain tissues of reciprocal crosses between B6 and JF1 mice[27, 28] (BJ: B6 mother × JF1 father; JB: JF1 mother × B6 father). These subspecies were selected because they harbor numerous polymorphic loci that facilitate allele-specific analyses. RT-PCR was conducted in reciprocal crosses (BJ and JB offspring), and imprinting status was assessed by direct sequencing of the amplified PCR products. The results showed that the mouse *Adam23* gene was biallelically expressed (allele ratio: 1.0–1.40), in contrast to the paternal expression observed in humans and ferrets (Figure 1c). These findings indicate that *Adam23* imprinting is species-specific and may be associated with both the presence of a gyrencephalic brain structure and the regulation of higher brain functions.

### Uniparental Expression of the Human *ADAM23* Gene upon Neural Differentiation

Given the species-dependent differences observed between ferrets and mice, and the prior report of paternal expression of human *ADAM23* in blood cells, we next examined the imprinting status of *ADAM23* in human neural cells. A previous study by Jadhav et al. described paternal expression of *ADAM23* in human blood [20]. However, in our panel of lymphoblastoid cell lines derived from healthy individuals, we did not identify an informative heterozygous polymorphism within *ADAM23*; therefore, allele-specific expression could not be assessed in these samples. Because of this limitation, we therefore assessed allele-specific expression in the neuroblastoma cell line SH-SY5Y, which constitutively expresses *ADAM23* and provides a well-established model for neuronal differentiation [27]. Sequencing of genomic DNA revealed a C/T polymorphism in the 3′ UTR of *ADAM23* (Figure 1d), enabling allele-specific expression analysis. RT-PCR and direct sequencing demonstrated biallelic expression in undifferentiated SH-SY5Y cells (allele ratio: 1.29). Following 18 days of neuronal differentiation induced by BDNF, db-cAMP, and retinoic acid, the same assay revealed a shift toward uniparental expression (allele ratio: 2.48; Figure 1d). Although the parental origin of the predominant allele could not be established, these findings indicate that *ADAM23* undergoes allele-specific regulation during neuronal differentiation.

Taken together, these results suggest that *ADAM23* may contribute to regulatory processes underlying higher brain functions, a possibility further explored in the Discussion.

### CpG Island Methylation Analysis of the Ferret *Adam23* Gene

Given that *Adam23* exhibits clear paternal expression in ferrets, we next sought to investigate the regulatory mechanism underlying this imprinting. Because promoter-associated CpG island methylation is a canonical mechanism of imprinting control, we examined DNA methylation within the upstream region of the ferret *Adam23* gene. To investigate whether CpG island methylation contributes to the imprinting of the ferret *Adam23* gene, we analyzed the upstream region of the gene. In this region, a gap was identified in the reference genome sequence (Ferret Genome Sequencing Consortium, 2011) (Supplementary Figure 5a) [40]. To obtain sequence information for this region and to facilitate analysis of potential regulatory elements controlling *Adam23* expression, we performed PCR amplification and sequencing (Supplementary Figure 5b). This approach successfully resolved the gap, yielding a 414 bp sequence that had previously been uncharacterized (Supplementary Figure 5c). Resolving this gap allowed us to identify previously undetectable features within the region, and subsequent computational analysis with MethPrimer [41] revealed two CpG islands (651 bp and 122 bp) within the upstream region (Supplementary Figure 5d).

We next assessed methylation of these CpG islands by bisulfite sequencing of a 1 kb region encompassing both islands. The region was divided into four segments (I∼IV), amplified from bisulfite-treated DNA, and sequenced after TA cloning of PCR products. No methylation was detected in any segment (Figure 1e), suggesting that paternal expression of ferret *Adam23* is not regulated by allele-specific methylation at the promoter region.

### Mouse *Adam23* Gene Knockdown and Neurite Outgrowth Enhancement

Because *Adam23* is paternally expressed in ferrets and therefore present at a functionally reduced dosage compared with mice, we next examined how reduced *Adam23* expression influences neuronal phenotypes by generating *Adam23* knockdown cells in a murine background. Knockdown experiments were performed in Neuro-2aTG cells [29], a murine neuroblastoma line that displays non-imprinted *Adam23* expression. Neuro-2aTG cells were transfected with an shRNA vector targeting *Adam23*, and stable knockdown lines were established. RT-PCR confirmed a 45–70% reduction in *Adam23* expression (Figure 3a). Cell proliferation assays showed no difference between knockdown and control cells (Figure 3b). In contrast, *Adam23* knockdown cells exhibited a pronounced increase in neurite outgrowth (Figure 3c), suggesting that Adam23 normally suppresses neurite elongation in this context.

**Figure 3.**
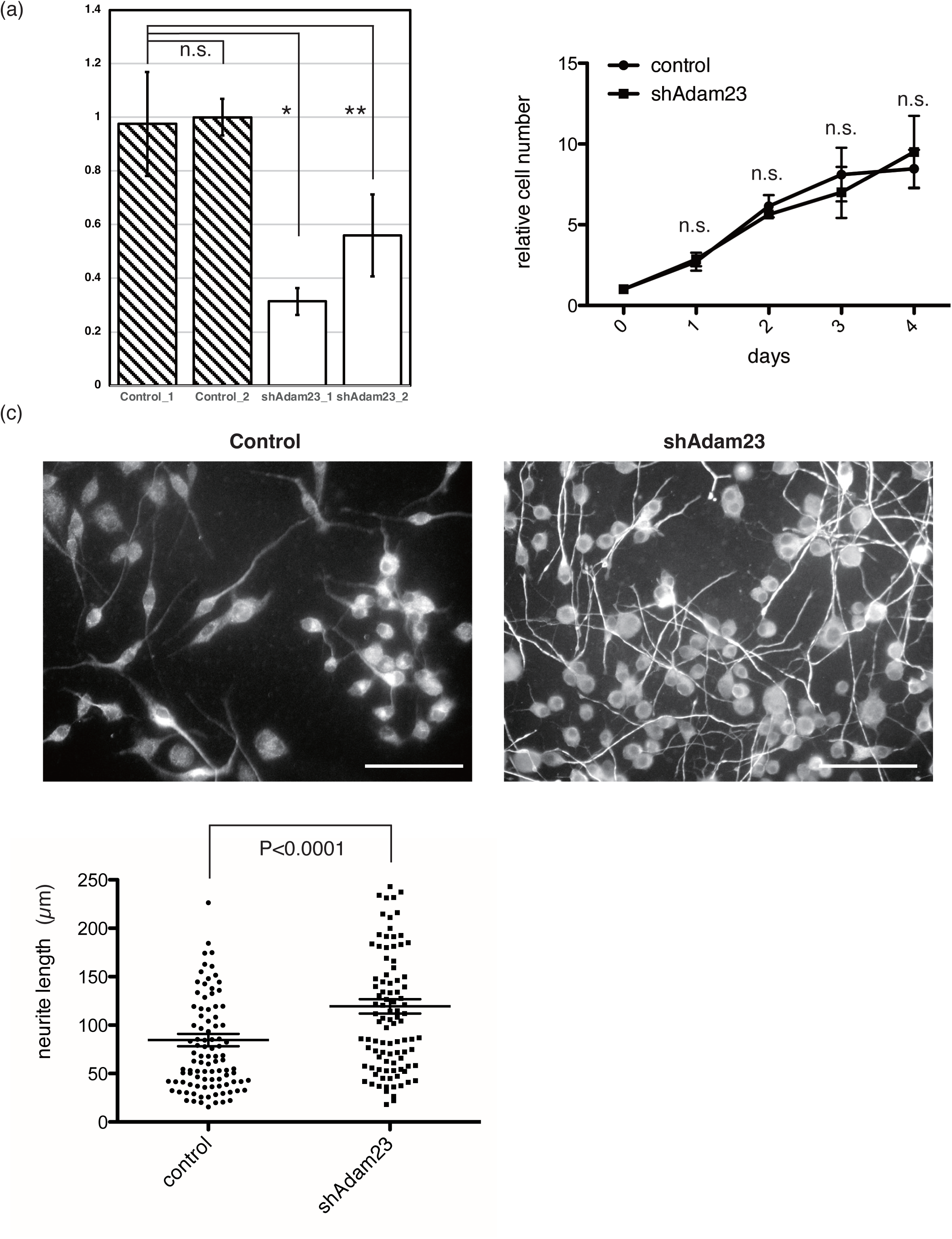
Knockdown of the mouse *Adam23* gene positively regulates neurite outgrowth. (a) Quantitative real-time PCR analysis of mouse *Adam23*. n.s., not significant; *p < 0.0001; **p < 0. 001. Two of the three established knockdown clones were selected for assessment of *Adam23* knockdown efficiency. (b) Cell proliferation curves of Neuro-2aTG cells following knockdown of mouse *Adam23*. n.s., not significant. Cell proliferation was evaluated using three independently established *Adam23*-knockdown clones. (c) Neuronal differentiation of *Adam23*-knockdown cells. Immunostaining was performed using an anti–βIII-tubulin antibody, and neurite length was quantified (n = 100). Scale bar, 100 μm. Neurite outgrowth analysis was performed using three independently established knockdown clones.

### Bovine *Adam23* Gene Overexpression and Cell Proliferation Enhancement

Because *Adam23* knockdown revealed dosage-sensitive effects on neurite morphology, we next asked whether increased *Adam23* dosage would exert reciprocal or distinct functional outcomes. To address this, we performed overexpression experiments in Neuro-2aTG cells. Stable Neuro-2aTG cell lines were generated by transfection with a plasmid carrying bovine *Adam23* cDNA under the CAG promoter, followed by drug selection. We chose the bovine gene so that transcripts derived from the transgene could be distinguished from endogenous mouse *Adam23* in PCR analyses. The bovine and mouse Adam23 proteins share 91% sequence identity (NP_001076927.1 and BAA83381.1, respectively), supporting the functional relevance of the bovine protein in this system. RT-PCR confirmed robust bovine *Adam23* expression in all established lines (Figure 4a).

**Figure 4.**
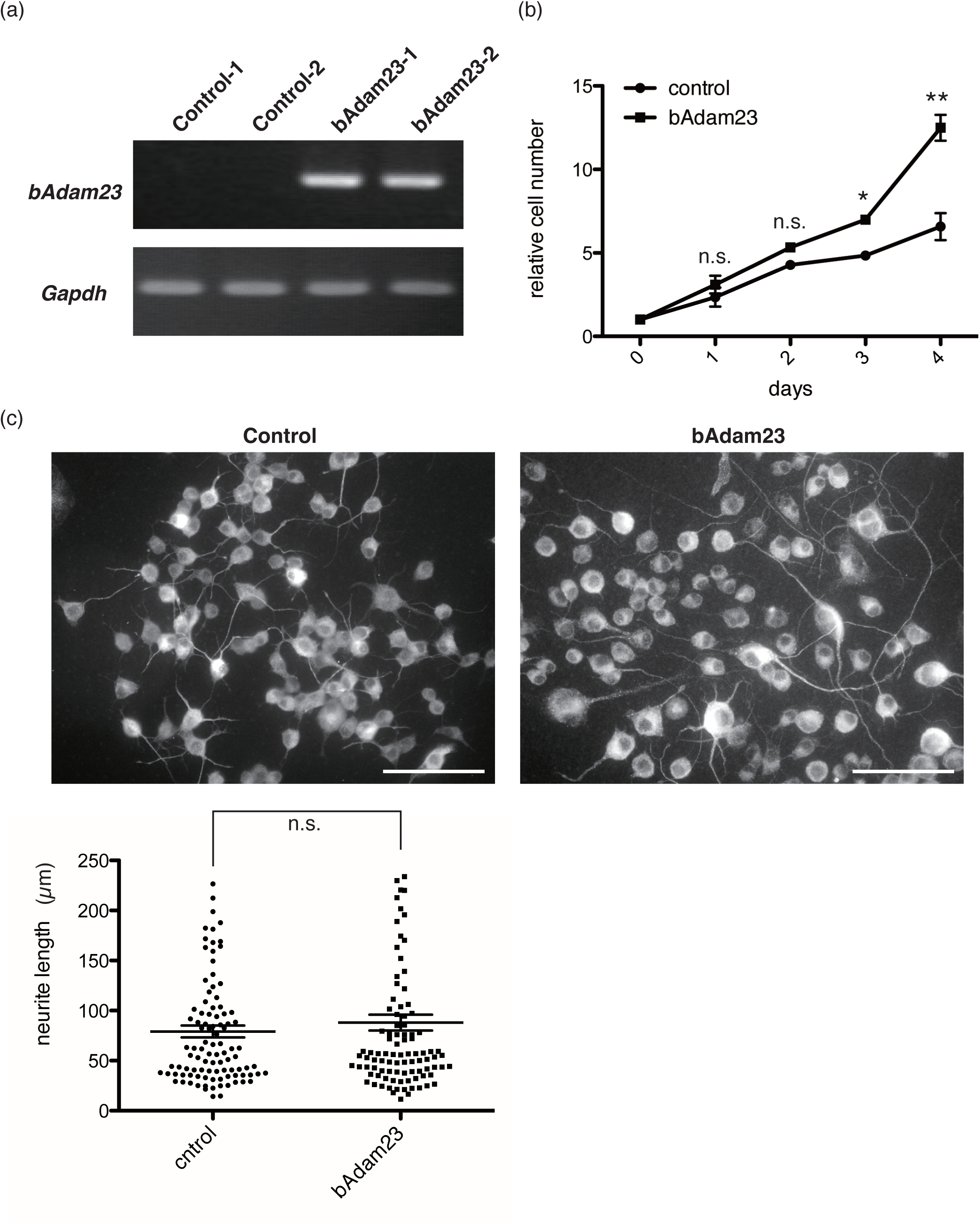
Overexpression of bovine *Adam23* positively regulates cell proliferation. (a) RT-PCR analysis of bovine *Adam23* expression in Neuro-2aTG cells transfected with a vector carrying bovine *Adam23* cDNA. Mouse *Gapdh* was used as a control. (b) Cell proliferation curves of cells overexpressing bovine *Adam23*. p < 0.05; p < 0.0001; n.s., not significant. Cell proliferation was evaluated using three independently established stable *Adam23*-overexpressing clones. (c) Neuronal differentiation of *Adam23*-overexpressing cells. Immunostaining was performed using an anti–βIII-tubulin antibody, and neurite length was quantified (n = 100). Scale bar, 100 μm. n.s., not significant. Neurite outgrowth analysis was performed using three independently established stable *Adam23*-overexpressing clones.

Functional assays demonstrated that *Adam23*-overexpressing cells exhibited significantly increased proliferation compared with control cells (Figure 4b). By contrast, neurite outgrowth did not differ significantly between overexpressing and control cells (Figure 4c). Thus, in Neuro-2aTG cells, elevated Adam23 dosage promotes proliferative capacity but does not influence neurite elongation.

## Discussion

Genomic imprinting has been extensively characterized in mice, yet its functional contribution to brain evolution and higher-order neural processes remains poorly understood[42–44]. The limited genetic diversity of inbred mouse models and the scarcity of imprinting studies in gyrencephalic species have restricted insights into how imprinting may influence the development of complex brain structures. In this study, we addressed this gap by systematically screening 65 candidate genes for imprinting in the ferret brain, a gyrencephalic mammal with cortical architecture more comparable to that of humans than mice.

Among the eight genes containing informative SNPs, *Pxdc1*, *Wrb* and *Ube3a* exhibited biallelic expression in the ferret brain despite being reported as imprinted in humans, suggesting that their imprinting may be primate-specific or restricted to certain tissues. In contrast, *Adam23*—a gene previously implicated in epilepsy and neurodevelopmental disorders [29,30]—showed robust paternal expression in ferrets. This imprinting was not associated with CpG island methylation, indicating that *Adam23* is regulated through a non-canonical imprinting mechanism. Such genes are frequently regulated by alternative mechanisms, including H3K27me3-mediated repression [31], raising the possibility that *Adam23* may be embedded within a broader epigenetically regulated chromosomal domain. Notably, the *Zdbf2*/*Gpr1* locus—located approximately 200 kb from *Adam23*—shows paternal expression in several mammalian species [45, 46] and may influence the imprinting state of *Adam23* through regional chromatin organization. Although we attempted to assess the imprinting status of *Zdbf2* and *Gpr1* in the ferret, no informative SNPs were identified, preventing allelic analysis. Future studies employing higher-resolution chromatin profiling and long-range epigenomic mapping in the ferret brain will be essential to determine whether *Adam23* is integrated into this imprinting domain and whether *Zdbf2/Gpr1* exerts cis-regulatory influence on *Adam23*.

A key finding of this study is the species divergence in *Adam23* imprinting: the gene is paternally expressed in gyrencephalic species (ferrets and humans) but biallelically expressed in lissencephalic mice. Moreover, within the ferret brain, paternal expression was most pronounced in the occipital cortex—a region with highly developed gyral architecture—suggesting that *Adam23* imprinting may play a region-specific role in cortical surface expansion. This pattern raises the intriguing possibility that *Adam23* imprinting contributes to the evolution of cortical complexity. Functional assays in Neuro-2aTG cells support this idea: *Adam23* overexpression increased cell proliferation, consistent with a role in neural stem cell expansion, whereas knockdown enhanced neurite outgrowth, potentially promoting the formation of more elaborate neuronal networks. Notably, our findings differ from those reported in human neural progenitor cells, where *ADAM23* overexpression promotes neuronal differentiation and neurite elongation, and its knockdown impairs these processes [37]. This apparent discrepancy likely reflects species-, lineage-, or developmental-stage–specific functions of ADAM23. Neuro-2aTG cells represent a transformed murine neuroblastoma line, whereas human NPCs correspond to early neural precursors with distinct transcriptional and epigenetic landscapes. Thus, ADAM23 may exert context-dependent, and even opposing, effects on neuronal maturation versus progenitor maintenance. This context dependence provides a plausible explanation for why Adam23 loss enhances neurite outgrowth in Neuro-2aTG cells, whereas its overexpression promotes differentiation in human NPCs.

Such dosage-sensitive and context-dependent functions are a hallmark of imprinted genes, which often regulate proliferation–differentiation balance in a cell-type–specific manner.

These observations align with the conflict hypothesis, which predicts that paternally expressed genes promote growth and resource acquisition. We propose that the shift from biallelic to paternal expression of *Adam23* may have provided an evolutionary advantage by modulating the balance between neural progenitor expansion and neuronal connectivity, thereby contributing to the emergence of gyrencephalic cortical structures. The fact that humans and ferrets share the same imprinting state for *Adam23*, despite substantial evolutionary distance, underscores the possibility that imprinting at this locus has been selectively maintained in species with higher cortical complexity.

In conclusion, our findings identify *Adam23* as a non-canonical imprinted gene in the ferret brain and suggest that its dosage contributes to neural stem cell dynamics and neurite development. These results highlight genomic imprinting not merely as a developmental curiosity but as a potential driver of mammalian brain evolution, and underscore the need for comparative imprinting studies across diverse taxa to fully elucidate how parental genomes shape the development and function of the mammalian brain.

## Acknowledgements

We thank Kotaro Nakamae for assistance with sequencing analyses of imprinted genes. This research was supported in part by JSPS KAKENHI Grant Numbers 22K06896, 20K07307, 23H00389, Japan Agency for Medical Research and Development (AMED) (JP24wm0625112), and The Mitani Foundation for Research and Development. These results were obtained from research (No. 23001) commissioned by the National Institute of Information and Communications Technology (NICT), Japan.

## Author Contributions

M.M-H. and S.H. conceived of and designed the study. M.M-H. and M.S. performed the experiments. K.S., Y.S., and H.K. contributed to the dissection and preparation of ferret brain tissues. M.M-H. wrote the manuscript and S.H. edited the manuscript. All authors approved of the final version of the manuscript.

---

**Supplementary Figure 1. Workflow for identifying imprinted genes in the ferret brain.**

(a) Based on the reports by Court *et al.* (Genome Res. 24, 554–569, 2014) and Jadhav *et al.* (BMC Biol. 17, 50, 2019), 65 candidate genes were selected for analysis.

Polymorphism analysis was conducted primarily on the 5′UTR and 3′UTR regions of these genes, and informative polymorphic sites suitable for imprinting analysis were identified in eight genes.

Among these eight genes, expression in the ferret brain was examined, and six genes were confirmed to be expressed.

Imprinting analysis was subsequently performed on these six genes, resulting in the identification of parent-of-origin–specific expression for three genes (*Adam23*, *Nhp2l1*, and *Atp10a*).

(b) The allelic ratio was calculated as the ratio of paternal to maternal peak heights in sequencing chromatograms.

(c) Paternal peak height of sequencing trace / maternal peak height of sequencing trace was determined for both cDNA and gDNA, and deviations in the cDNA/gDNA ratios were used to assess imprinting status.

**Supplementary Figure 2. *Fam50b* and *Ptchd3* were not detectably expressed in the ferret brain.**

RT-PCR analysis of *Fam50b* and *Ptchd3* expression in ferret tissues:

1. olfactory bulb, 2. frontal cortex, 3. parietal cortex, 4. occipital cortex, and 5. liver.

Ferret *Gapdh* served as a control.

**Supplementary Figure 3. Results of imprinting analysis for three candidate genes in the ferret brain.**

(a) The ferret *Pxdc1* gene exhibits biallelic expression. Arrowheads indicate polymorphic sites used for imprinting analysis. Allelic ratios calculated for imprinting analysis are shown beneath the cDNA chromatograms. gDNA, genomic DNA; cDNA, complementary DNA. Pup1 and Pup2 represent offspring derived from the parental combinations shown on the left (Father × Mother). No informative SNPs were identified in Pup2 for imprinting analysis.

(b) The ferret *Wrb* gene displays biallelic expression. Arrowheads indicate polymorphic sites used for imprinting analysis. Allelic ratios calculated for imprinting analysis are shown beneath the cDNA chromatograms. Pup1 and Pup2 represent offspring derived from the parental combinations shown on the left (Father × Mother).

(c) The ferret *Ube3a* gene displays biallelic expression. Arrowheads indicate polymorphic sites used for imprinting analysis. Allelic ratios calculated for imprinting analysis are shown beneath the cDNA chromatograms. Pup1 and Pup2 represent offspring derived from the parental combinations shown on the left (Father × Mother).

**Supplementary Figure 4. The ferret *Atp10a* gene shows predominantly maternal expression** The ferret *Atp10a* gene shows predominantly maternal expression. Arrowheads indicate polymorphic sites used for imprinting analysis. Allelic ratios calculated for imprinting analysis are shown beneath the cDNA chromatograms. Pup1 and Pup2 represent offspring derived from the parental combinations shown on the left (Father × Mother).

**Supplementary Figure 5. Sequencing of the upstream region of the ferret *Adam23* gene and identification of a CpG island.**

(a) Upstream genomic region of the ferret *Adam23* gene (GL897001: 5,251,501–5,253,500). Arrowheads indicate PCR primers used to amplify the genomic region containing the sequence gap.

(b) PCR amplification of an 880-bp DNA fragment spanning the genomic gap.

(c) Determination of the gap sequence. The newly identified gap sequence is shown in red.

(d) Identification of a CpG island using MethPrimer (https://www.urogene.org/cgi-bin/methprimer/methprimer.cgi).

